# The *Helicobacter pylori* orphan ATTAAT-specific methyltransferase M.Hpy99XIX plays a central role in the coordinated regulation of genes involved in iron metabolism

**DOI:** 10.1101/2024.11.27.625509

**Authors:** Wilhelm Gottschall, Florent Ailloud, Christine Josenhans, Sebastian Suerbaum

## Abstract

*H. pylori* genomes contain a large and variable portfolio of methyltransferases (MTases), creating a highly diverse methylome. Here, we characterize a highly conserved ATTAAT-specific MTase, M.Hpy99XIX, the only *H. pylori* MTase never associated with an endonuclease (“orphan” MTase). Inactivation of M.Hpy99XIX resulted in a significant change in the transcription of >100 genes, despite the fact that only a small subset of their promoter regions contained an ATTAAT target motif. Patterns of transcriptional change showed significant correlations with changes reported for *H. pylori* mutants in the ArsRS regulators involved in iron regulation. The MTase inactivation also caused a higher susceptibility to diverse metal ions as well as iron chelation and oxidative stress. These phenotypes could be traced back to the methylation of single motifs in the promoter regions of iron transporters *frpB1* and *fecA1*. Altogether, methylation of individual motifs in promoters can have a large downstream effect causing major changes to metabolic pathways. These findings suggest that the methylome represents a universal and dynamic interface connecting genome diversity and transcriptional regulation. Very recently, a new ecospecies *Hardy* of *H. pylori* has been reported. M.Hpy99XIX is present in the majority of “normal” (*Ubiquitous*) *H. pylori* strains, whereas no single *Hardy* strain contained this gene, consistent with other reported differences between *Hardy* and *Ubiquitous* strains related to iron/metal homeostasis. ATTAAT methylation is intricately connected with the bacterial transcriptional network, highlighting the important role of bacterial epigenetic modifications in bacterial physiology and pathogenesis.

## Introduction

*Helicobacter pylori* is a genetically highly diverse bacterial species that can colonize the stomach mucosa of humans for many decades and can lead to severe diseases such as peptic ulcers, gastric adenocarcinoma and lymphoma of the mucosa-associated lymphoid tissue (MALT) [1–3]. *H. pylori* has a large repertoire of Restriction-Modification (RM) systems [4–7]. While the pangenome of *H. pylori* comprises more than 100 methyltransferases (MTases), only a minority of these is highly conserved across the diverse phylogeographical populations within the species [8]. Each strain carries a specific repertoire of RM systems, acting on a unique pattern of target motifs in the genome, resulting in virtually unique methylomes [8, 9]. In the last decade, research on DNA methylation and its role in bacterial physiology has been accelerated by the development of novel technologies which permit to detect methylation at single base resolution (in particular, Single Molecule Real-Time sequencing [10], and, more recently, Oxford Nanopore sequencing [11]). For example, methylation of the motif GCGC by the MTase M.Hpy99III, one of the two MTases present in all *H. pylori* strains, is associated with multiple important phenotypes, such as adhesion to gastric cells, natural competence, bacterial cell shape or copper tolerance [12]. Similarly, three other, less conserved MTases have also been connected to important traits such as motility and tolerance to oxidative stress [13].

The phenotypic effects connected to methylation in *H. pylori* have, in some cases, been shown to be mediated by changes in gene expression [12–15]. However, the connection between methylation and transcription appears to be highly complex and significant variation was observed among different MTases and between strains of *H. pylori* [12, 13]. For the M.Hpy99III MTase, transcriptional regulation is based on the methylation of GCGC motifs within promoter regions [12]. This has been demonstrated for four distinct genes using targeted mutagenesis and reporter assays [12, 16]. Because genes carrying a target motif within their promoter generally account only for a small proportion of genes regulated by MTases [12–15], additional regulatory mechanisms and large downstream transcriptional effects are likely at play.

In bacterial genomes, MTase genes are frequently coupled with genes coding for a cognate endonuclease, recognizing the same sequence motif as the MTase, constituting RM systems. These are thought to be primarily involved in defense against phages and invading foreign DNA [17, 18]. In type II RM systems, the endonuclease is not essential for the activity of the MTase and therefore type II MTase genes can persist independently as so-called “orphan” genes [19]. Since selection pressures acting on orphan MTases are not influenced by those acting on a cognate endonuclease anymore, orphan MTases constitute an ideal model to better understand the mechanisms connecting methylation, transcription and phenotypic effects.

Here, we determined that M.Hpy99XIX is the only strictly orphan MTase in *H. pylori* and that its target motif ATTAAT is enriched in promoter regions. A large group of genes was differentially regulated after inactivation of ATTAAT methylation, disrupting the iron homeostasis regulatory pathway on the transcriptomic and phenotypic levels. These phenotypic effects could be recapitulated by the targeted modification of a single ATTAAT motif located within the promoter region of an iron acquisition gene, demonstrating how DNA methylation can have large effects on transcriptional networks and core bacterial processes by only modulating a very limited number of genes directly. On a larger scale, this study substantiates the role of the methylome as an integrated layer of diversity in *H. pylori* inter-connected with the genome through individual sequence motifs and with the transcriptome via methylation-based regulation.

## Results

### 1. M.Hpy99XIX is the only strictly orphan MTase in *H. pylori*

Methyltransferases without a coupled endonuclease, frequently referred to as orphan MTases, cannot fulfill their presumed original function as part of RM systems, such as defense against invading DNA. In bacteria, orphan MTases have been frequently associated with transcriptional regulation in recent years. To determine the proportion of *H. pylori* MTases that are not part of canonical RM systems, we analyzed a collection of ∼400 geographically diverse genomes of *H. pylori* that was the basis of our previous characterization of the methylome diversity in *H. pylori* [8]. In particular, we determined the frequency of all known *H. pylori* type II MTase and cognate endonuclease genes in this genome collection [20]. When no cognate endonuclease could be found, the genes neighboring the MTase locus were examined to confirm that they were not homologs of endonucleases.

Remarkably, we could only identify a single MTase gene, M.Hpy99XIX, that was never associated with an endonuclease across the whole species (Fig. 1A). The M.Hpy99XIX MTase is highly conserved and predicted to be active in ∼90% of strains in the collection. We also analyzed the recently released HpGP collection of 1012 finished *H. pylori* genomes [21]. In this collection, the M.Hpy99XIX MTase was predicted to be functional in ∼94% of the isolates and found exclusively at a single locus, between the genes *hopJ* (*jhp0429*), coding for an outer membrane protein, and *hepT* (*jhp0431*), encoding a DD-heptosyltransferase (Fig. 1B). Subsequent analysis showed that the MTase gene was fragmented in 43 of the 1012 strains (4.3%) due to nonsense mutations, which likely render it inactive. In a smaller subset of only 10 strains (1%), the MTase gene was completely absent with no apparent remnants or excision site. While the strains affected by nonsense mutations were not obviously genetically related, the strains lacking the M.Hpy99XIX gene all belonged to a single *H. pylori* subpopulation, hspIndigenousAmerica.

**Figure 1.**
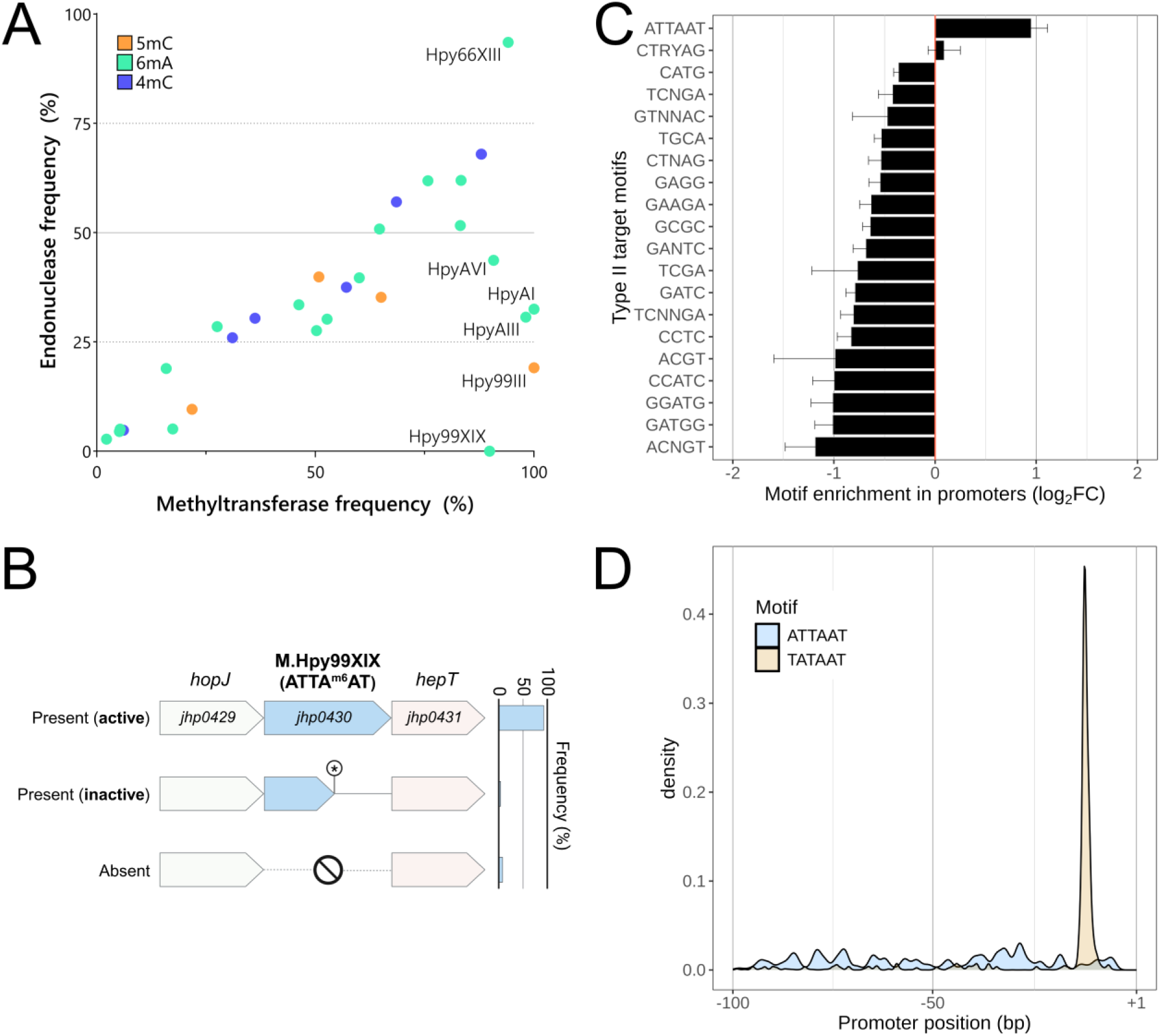
Genetic characterization of the *H. pylori* M.Hpy99XIX methyltransferase and its target motif ATTAAT. **A.** Frequencies of 26 type II *H. pylori* methyltransferase (x-axis) and cognate endonuclease (y-axis) genes in a collection of geographically diverse 398 *H. pylori* genomes. Frequencies were normalized across four geographical populations of *H. pylori* (hpAfrica1, hpAfrica2, hpEurope, hpAsia2) **B.** Genetic context and frequency of the complete, inactivated and empty locus in a distinct collection of 1029 *H. pylori* genomes. The complete form is by far the most frequent and is always flanked by the *hopJ (jhp0429)* and *hepT (jhp0431)* genes. **C.** Enrichment analysis of type II motifs performed by calculating the ratio of motif frequencies in promoter regions and in the whole *H. pylori* genome. Log_2_ fold-changes > 1 and < 1 indicate motifs more and less frequent in promoters compared to the rest of the genome, respectively. Error bars indicate the standard-deviation across all *H. pylori* genomes analyzed. The first 20 motifs with the highest absolute Log_2_ fold-changes are shown. **D.** Positional bias analysis of the ATTAAT target motif (blue) and the TATA box (orange) within promoter regions. The ATTAAT motif does not appear at any preferred position.

### 2. The ATTAAT motif is the only MTase target sequence enriched in promoter regions in *H. pylori*

The M.Hpy99XIX MTase targets the palindromic motif ATTAAT and catalyzes the N6-methyladenosine modification at the 5^th^ position. In *H. pylori* and other bacterial species, methylation of single motifs located within promoter regions has been associated with regulation of the downstream transcriptional unit [12, 16]. Compared to some other motifs modified by type II enzymes, the ATTAAT motif is relatively infrequent in the *H. pylori* genome with an average of 553 motifs per megabase [8] and thus is not likely to be randomly observed near promoter regions. Consequently, we determined how often this motif appears in promoter regions and thus potentially directly acts on downstream gene expression.

We calculated ratios (as fold change FC) between the relative motif frequencies within promoters and the corresponding motif frequencies in the complete genome, for the ATTAAT motif and selected other known type II motifs (Fig. 1C). Consequently, this analysis describes whether a given motif is either over- (log_2_FC > 0) or under- (log_2_FC < 0) represented within promoter regions. Almost all type II motifs were significantly underrepresented in promoter regions whereas the ATTAAT motif was significantly enriched with a positive log_2_FC of 0.98, signifying that it was observed almost twice as frequently in promoter regions compared to the rest of the genome. On average, the ATTAAT motif was observed in 37±5 promoter regions per strain, representing approximately 5% of the promoter regions in *H. pylori*.

Regulatory motifs are often located in fixed positions relative to the transcription start sites. Based on all promoters carrying ATTAAT motifs, we determined the probability distribution function of the motif position and of the Pribnow box (TATAAT) typically located at -10 as a reference [22]. In contrast to TATAAT, no specific positions were favored by the ATTAAT motif within the promoter region (Fig. 1D).

### 3. ATTAAT methylation by M.Hpy99XIX has a large effect on the *H. pylori* transcriptome

In order to determine whether M.Hpy99XIX and the methylation of ATTAAT have a role in gene expression, we compared the transcriptomes of wild-type *H. pylori* strain J99 [23], an isogenic M.Hpy99XIX knockout (KO) mutant, and a functionally complemented strain, using RNA-Seq analysis (Table S2). Differential gene expression analysis revealed that 85 genes were downregulated in the ATTA^m6^AT-methylation deficient strain compared to the wild-type (Log_2_fold-change < 1 and FDR adjusted p-value <0.01) whereas 28 genes were upregulated (Log_2_fold-change > 1 and FDR adjusted p-value <0.01) (Fig. 2A), altogether representing ∼8% of the total genes in the *H. pylori* J99 genome. The functional complementation (Fig. S1) restored the expression of 81.2% of downregulated and of 96.6% of upregulated genes to their wild-type expression levels. Ten and five new genes were down- and upregulated in the complemented strain compared to the wild-type, respectively, possibly owing to the increased expression of the MTase in the complemented strain which is a consequence of the insertion of the intact MTase gene in the highly transcribed urease gene cluster (Fig. S2).

**Figure 2.**
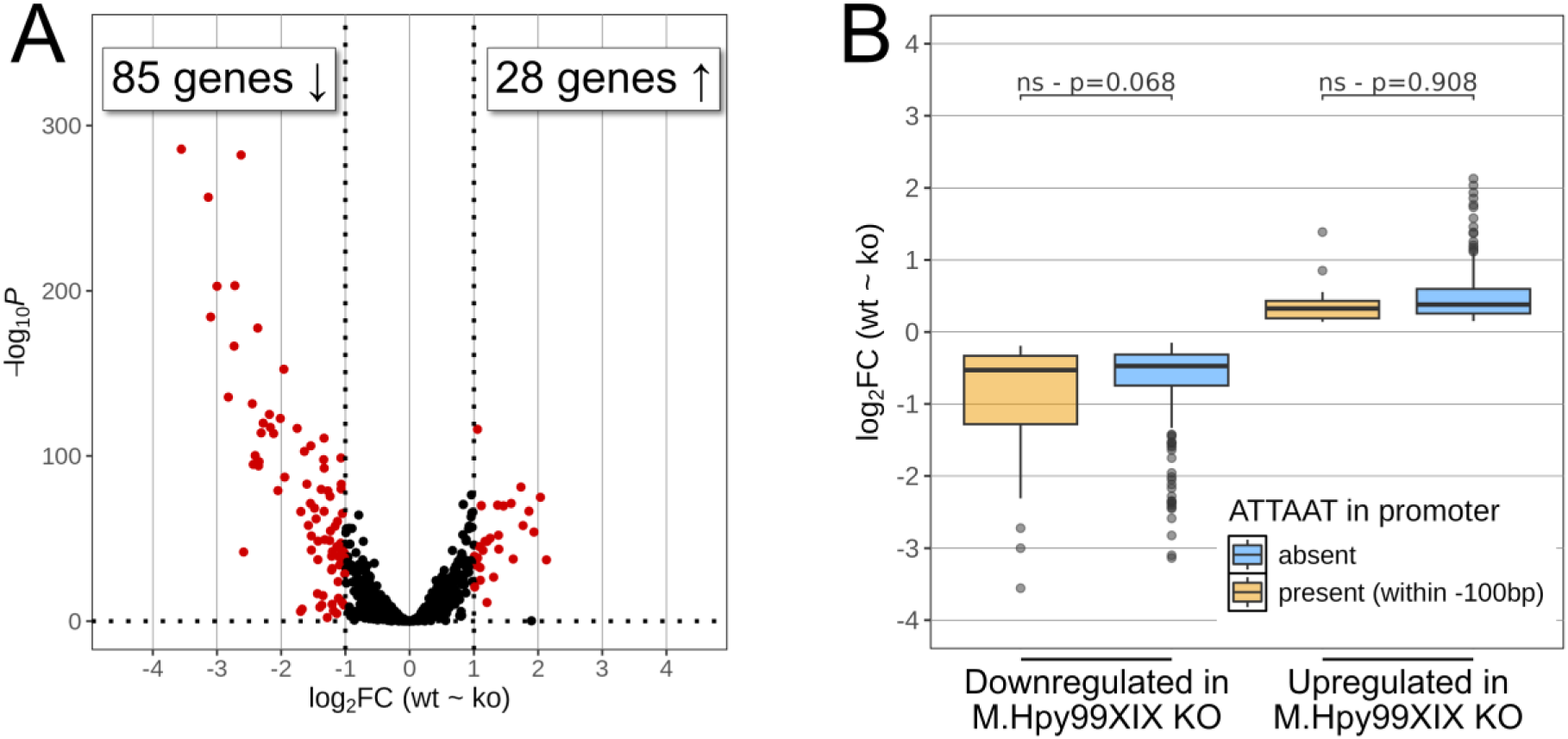
Effects of M.Hpy99XIX on the transcriptome of *H. pylori* strain J99. **A.** Volcano plot of *H. pylori* J99 wild-type compared to *H. pylori* J99 M.Hpy99XIX knockout strain. A negative and positive log_2_ fold change indicate a gene downregulated (left) or upregulated (right) in the knockout strain compared to the wild-type, respectively. Genes with an adjusted p-value < 0.01 and an absolute log_2_ fold change > 1 are shown in red. **B.** The log_2_ fold changes of down- (left) and upregulated (right) genes were compared (Student’s t-test) according to presence (orange) or absence (blue) of an ATTAAT motif within 100 bp of their transcription start sites (promoter regions).

Next, we asked whether the differential gene expression resulting from the absence of ATTA^m6^AT methylation could be correlated with the position and local frequency of ATTAAT motifs across the genome. We calculated a motif enrichment score for ATTAAT in coding sequences (see Methods). An average of 0.3±0.6 motifs was observed per CDS and 77 CDS showed a significant enrichment. Nevertheless, no correlation was found between enrichment score and gene expression (Fig. S3, Table S3). Next, we turned to motifs residing in promoter regions. An ATTAAT motif could be detected in the promoter region of 8% and 3% of up- and downregulated genes, respectively (Table S3). While no significant association was found (p=0.068), downregulated genes that were linked to an ATTAAT motif in a promoter region-leaned toward lower fold-changes than other downregulated genes (Fig. 2B). No similar trend was observed for upregulated genes (p=0.908). Furthermore, ATTAAT motifs located in promoter regions of differentially expressed genes were not located at a specific position relative to the transcriptional start site, yet 5 out of these 8 motifs were found in the range of 67 to 80 bp upstream of the TSS (Fig. S4). Altogether, the data suggest that the majority of genes in our transcriptomic dataset are not directly controlled by ATTA^m6^AT methylation.

### 4. ATTA^m6^AT-based regulation is amplified by core transcriptional regulators

The regulatory pathways of *H. pylori* have been extensively characterized, and core transcriptional regulators have been described [24]. Therefore, we asked whether significant overlaps existed between known *H. pylori* regulons and genes regulated through ATTA^m6^AT methylation. Interestingly, we were able to find a significant association between genes regulated by ATTAAT methylation and four out of the eight known regulon datasets we tested and associated with acid and iron-response as well as the ArsRS, HspR, HrcA, NikR, HP1021 and CrdRS transcriptional regulators [25–30] (Table S3).

Transcriptomic regulation connected to ATTA^m6^AT methylation mirrored the changes associated with the ArsRS two-component system (Fig. S5A) [25] and were opposite to the ones that have been linked to the HspR regulator (Fig. S5B) [26]. 66% (21/32) of the genes downregulated in the ArsRS KO were also significantly downregulated in the M.Hpy99XIX KO strain (FDR adjusted p-value < 0.01 and log_2_FC < 0) while 39% (27/70) of the genes upregulated in the ArsRS KO were significantly upregulated in the M.Hpy99XIX KO strain (FDR adjusted p-value < 0.01 and log_2_FC > 0). By contrast, 80% (8/10) of the genes downregulated in the HspR KO were upregulated in the M.Hpy99XIX KO strain while 80% (8/10) of the genes upregulated in the HspR KO were downregulated in the M.Hpy99XIX KO strain. When considering genes with at least a 50% difference in expression (equivalent to an absolute log_2_FC = 0.58 in the M.Hpy99XIX KO strain), a moderate correlation could be observed between the fold-changes of differentially expressed genes in the M.Hpy99XIX KO strain with the ArsRS regulon (ρ= 0.59, p < 0.001) but not with the HspR regulon (ρ= 0.48 p = 0.1)(Fig. S6). For instance, the ferric uptake regulator (Fur, *jhp0397*) was both downregulated in the absence of ATTA^m6^AT methylation and by ArsRS [25]. In contrast, the chaperone and stress-response proteins, GroES (*jhp0009*) and DnaK (*jhp0101*) were upregulated in the M.Hpy99XIX KO strain whereas they were downregulated by HspR.

Furthermore, ATTA^m6^AT-based regulation also matched transcriptional changes associated with regulatory responses to low pH which is largely dependent on ArsRS (Fig. 3A) [25] and extracellular iron content (Fig. 3B) [27]. Specifically, 59% (10/17) and 55% (17/31) of genes repressed and induced at low pH were up- and downregulated in the M.Hpy99XIX KO strain, respectively, while 66% (50/76) and 59% (29/49) of genes repressed and induced in response to a high iron concentration were up- and downregulated in the M.Hpy99XIX KO strain, respectively. Moreover, strong correlations could be observed between the fold-changes in the M.Hpy99XIX KO strain with the acid-responsive regulon (ρ= 0.85, p < 0.001) and the iron-responsive regulon (ρ= 0.70, p < 0.001)(Fig. S6).

**Figure 3.**
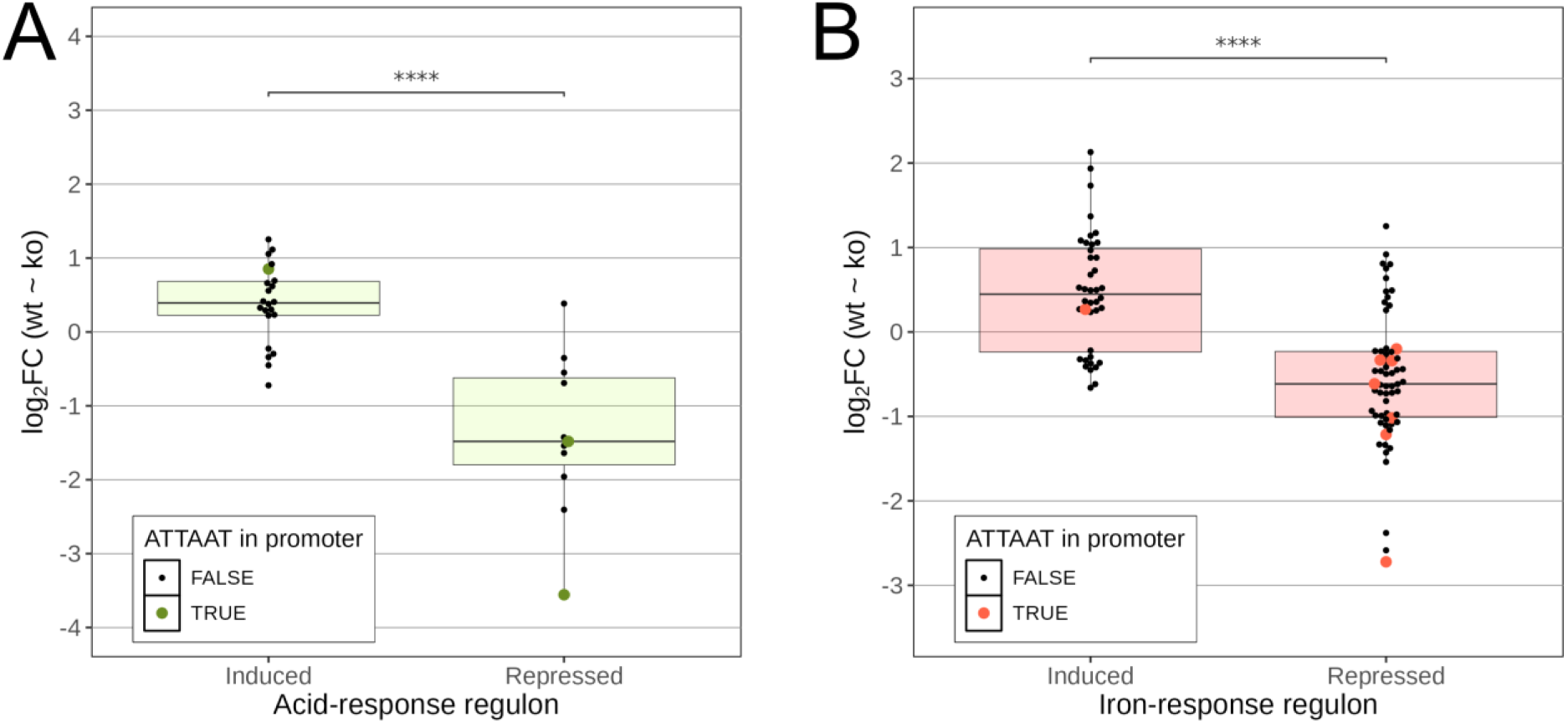
Comparison between M.Hpy99XIX transcriptional effects and environment responsive regulons. **A.** Comparison of RNA-Seq expression data of M.Hpy99XIX KO against WT with the acid-responsive regulon (Loh et al. 2021) split into induced and repressed genes. Student’s t-test: *** p<0.001, **** p<0.00001. **B.** Comparison of RNA-Seq expression data of M.Hpy99XIX KO against WT with the iron-responsive regulon (Vannini et al. 2024) split into induced and repressed genes. Student’s t-test: *** p<0.001, **** p<0.00001. Log_2_ fold-changes of significantly differentially expressed genes (adjusted p-value < 0.01) are represented by individual points and by a boxplot. The presence of an ATTAAT motif within 100 bp of the TSS of each gene is indicated by a larger coloured point.

These results reflect the inter-connection of these regulatory networks in *H.* pylori. For instance, acid adaptation has been connected to both the ArsRS [31] and Ferric uptake regulator (Fur) transcriptional regulators, despite Fur being initially associated with the control of iron homeostasis [27]. Collectively, this indicates that the ATTAAT transcriptional response may serve to fine-tune specific regulatory pathways existing in *H. pylori*.

### 5. Transcriptional induction of iron transporter genes by ATTAAT methylation in promoters

To further understand the connection between the ATTAAT transcriptional response and the major regulatory pathways, we looked for ATTAAT motifs within the promoter regions of genes that displayed a similar transcriptional behavior in the M.Hpy99XIX KO strain and in the acid- and iron-responsive regulons. ATTAAT motifs were observed in the promoter regions (within -100 bp of the TSS) in three and eight genes differentially expressed in both the M.Hpy99XIX KO strain and the acid-responsive and iron-responsive regulons, respectively. In both cases, ATTAAT motifs were observed more frequently in down-than upregulated genes, indicating that ATTA^m6^AT methylation in promoter regions generally leads to an increase of gene expression. In particular, seven genes downregulated in response to iron and in the M.Hpy99XIX KO strain carried an ATTAAT motif within their promoter region, including the two major iron transporter genes *fecA1* (*jhp0626*) and *frpB1* (*jhp0810*) [32]. This observation implies that the pleiotropic regulatory effects connected to the inactivation of the ATTAAT MTase might originate from a defect in iron transport and dysregulation of iron homeostasis.

Up to now, regulation of gene expression via methylation of promoter-bound target sequences has not been demonstrated for the ATTAAT motif. To this end, we generated an isogenic point mutant in which only the ATTAAT motif located in the promoter of the *frpB1* gene (at -80 bp) was mutated in order to prevent methylation at this specific location. Specifically, the methylatable adenine, in the fifth position of the motif, was changed to a thymine using targeted mutagenesis (Fig. 4A). Similar to the M.Hpy99XIX KO strain, the *frpB1* point mutant strain showed a significant decrease (>50%) in expression of the *frpB1* gene compared to the wild-type (Fig. 4B). This indicates that the expression of the *frpB1* gene is specifically and directly increased by the methylation of the ATTAAT motif within its promoter.

**Figure 4.**
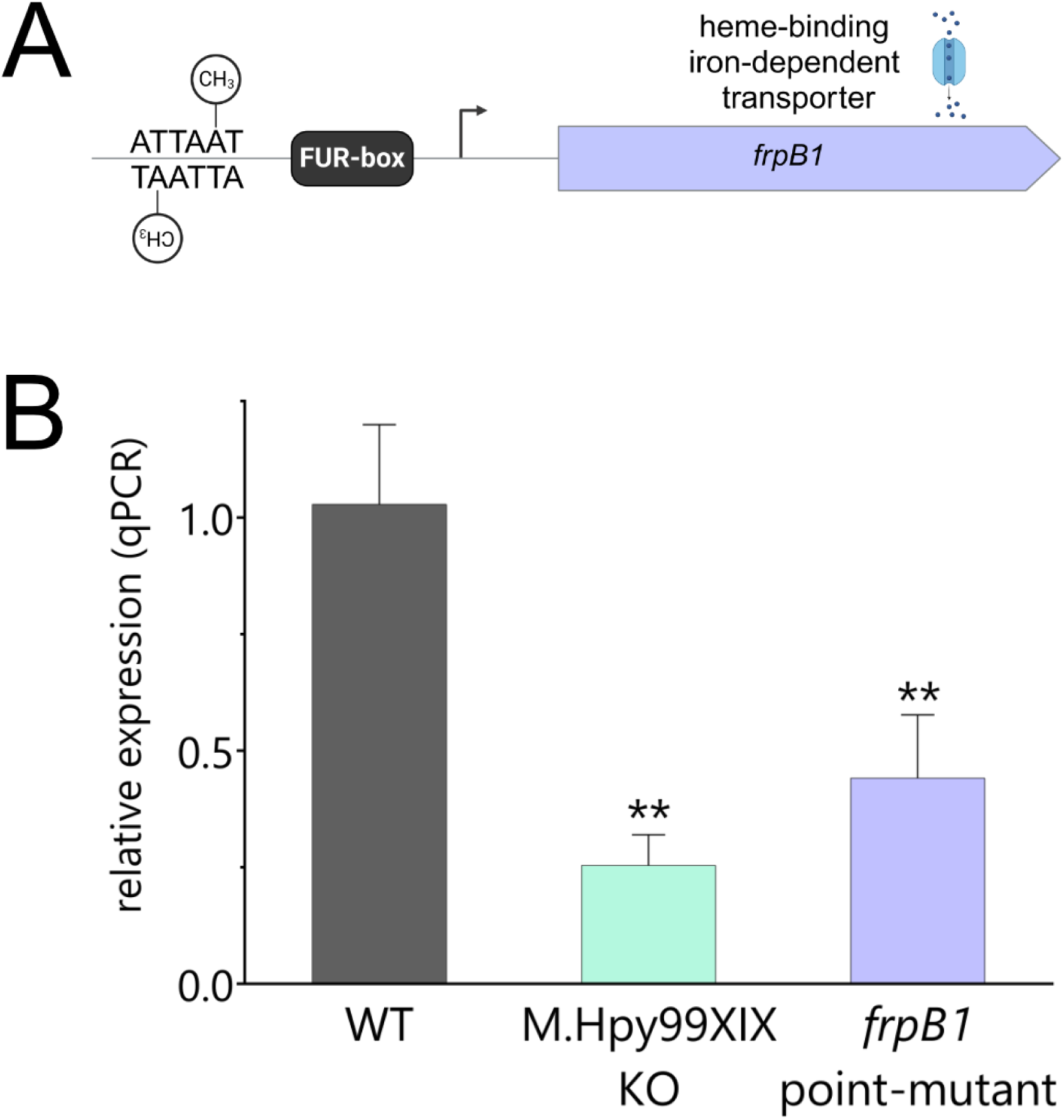
Characterization of the effect of ATTAAT methylation within the promoter region of *frpB1*. **A.** Genetic context of the *frpB1* gene, coding for a heme-binding iron-dependent transporter. Upstream of the transcription start site is i) a FUR-box targeted by the transcription factor Fur and ii) an ATTAAT motif. A point mutant strain was created in which the methylatable A within the motif was changed to a T via targeted mutagenesis in order to prevent methylation at this position. **B.** Comparison of *frpB1* expression in the M.Hpy99XIX knockout and the point mutant to the wild-type strain by qPCR. Student’s t-test: ** p<0.01. Error bars indicate the standard deviation of 2-5 replicates.

Next, we repeated the experiment with the second transporter gene, *fecA1*. Because *fecA1* contains two ATTAAT motifs in its promoter region (located at -67 and -96 bp of the *fecA1* TSS), we created two isogenic point mutants and determined that mutation in only one of the motifs (the one farther from the TSS) led to a lower expression of the *fecA1* gene (∼50%) (Fig. S7). By contrast, the mutant of the motif closer to the TSS showed a two-fold increase of *fecA1* expression compared to the wild-type. A double mutant of both ATTAAT motifs showed a four-fold increase of *fecA1* transcript, suggesting that gene regulation can be adjustable depending on the number, and position, of methylated motifs within promoter regions.

Critical regulatory elements are generally conserved across bacterial strains [33, 34]. We looked at the sequence diversity of the two ATTAAT motifs involved in the control of expression of the *frpB1* and *fecA1* genes and found these motifs to be highly conserved at 95% and 82% in our *H. pylori* genome collection, respectively. The motif leading to reduced expression of *fecA1* was also highly conserved at 97%. Compared to the average 8% conservation of ATTAAT motifs in the same *H. pylori* genome collection, this indicates that these promoter-bound motifs could be maintained by selection due to their role in regulating gene expression.

### 6. M.Hpy99XIX regulates metal homeostasis and oxidative stress

Because FrpB1 is involved in iron acquisition, a decreased expression could have a downstream effect on iron metabolism as well as on associated pathways and regulatory networks. Furthermore, since the *frpB1* gene only has a single ATTAAT motif in its promoter region, it represents a simplified model to understand the effects of ATTA^m6^AT methylation. To determine the extent of this theoretical effect, we tested the tolerance of the ATTA^m6^AT deficient strains to different conditions.

First, we compared the effects of supplementation with iron (1 mM (NH4)_2_Fe(SO_4_)_2_) to those of iron depletion (75 µM 2,2′-dipyridyl (dpp)). Compared to the *H. pylori* J99 wild-type strain, the isogenic M.Hpy99XIX KO strain and the *frpB1* point mutant showed a significant growth defect in both conditions (Fig. 5). Specifically, the KO and point mutant strains had a delay in growth leading them to grow 70% and 25% slower than the wild-type in iron supplemented media. Under standard culture conditions, no significant differences in growth could be observed between the wild-type, M.Hpy99XIX KO and *frpB1* point mutant strains (Fig. S8).

**Figure 5.**
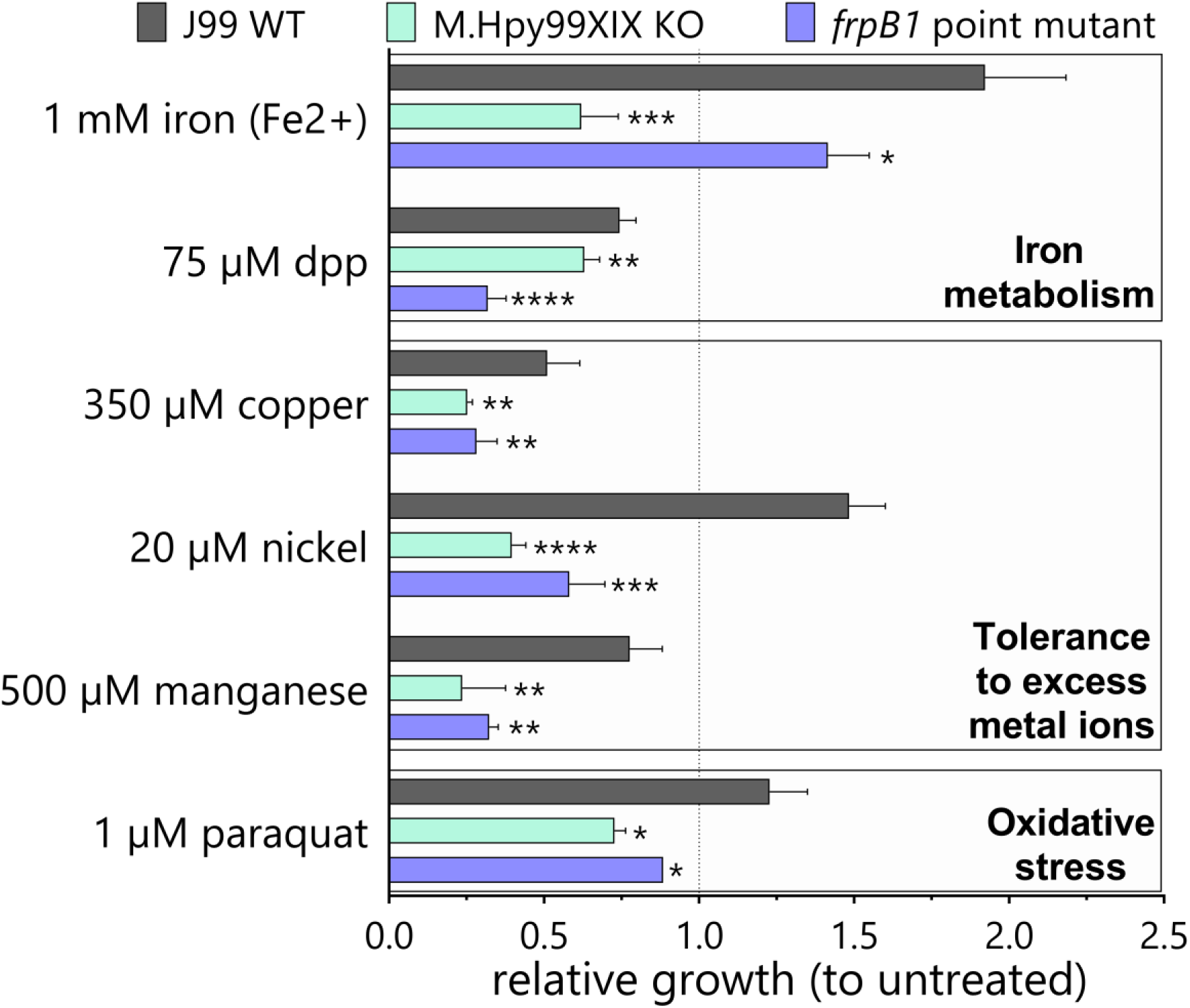
Impact of ATTAAT methylation on growth under environmental conditions related to iron metabolism. Growth of WT and mutants under iron, iron chelation (dpp), copper, nickel, manganese and oxidative stress (paraquat) conditions was normalised to growth without treatment and compared after 20 hours. Student’s t-test: * p<0.05, ** p<0.01, *** p<0.001, **** p<0.0001. Error bars indicate the standard deviation of 3-5 replicates.

Through inter-connected regulatory networks, iron metabolism has been shown to be influenced by other metal ions. Consequently, we repeated the growth assay in the presence of high concentrations of copper, nickel and manganese. Interestingly, all metal ions affected bacterial growth in both, M.Hpy99XIX KO and *frpB1* point mutant strains. Nickel had the strongest effect, causing a 75% and 40% slower growth in the KO and point mutant, respectively. In addition, the allosteric regulation of Fur has been shown to be triggered by reactive oxygen species. Following exposure to paraquat, we observed that both the M.Hpy99XIX KO and *frpB1* point mutant strains exhibited a significant growth defect, suggesting they are more susceptible to oxidative stress.

Altogether, in the two mutant strains (M.Hpy99XIX KO and *frpB1* point mutant), iron homeostasis thus appeared to be dysregulated in the presence of a high or low extracellular iron content that is most likely leading to toxicity and reduced metabolic activity, respectively.

### 7. Indirect transcriptional effects mediated by direct ATTA^m6^AT-based regulation of gene expression

The data shown in the previous sections demonstrate that the disruption of iron homeostasis observed in the ATTAAT methylation deficient strains is rooted in changes in the Fur regulatory network. In the M.Hpy99XIX KO strain, we determined that a significant proportion of the transcriptomic effect was similar to the changes observed in response to extracellular iron content. We selected six genes that were differentially regulated in the M.Hpy99XIX KO strain and known from previous studies to be involved in iron metabolism [27], and analyzed their expression in the *frpB1* point mutant strain. All six genes carry a Fur box in their promoter [27, 35]. Among the selected genes, two were upregulated in the M.Hpy99XIX KO strain, coding for Pfr, a ferritin protein involved in iron storage [36], and HydA, a subunit of a [Ni-Fe] hydrogenase [37]. In the *frpB1* point mutant, *hydA* had a similar trend but was not significantly upregulated whereas *pfr* showed a completely opposite expression pattern (Fig. 6). The remaining four genes were downregulated in the M.Hpy99XIX KO strain and code for FecA1, an iron transporter [32]; RibBA and PdxJ, enzymes involved in riboflavin [38] and vitamin B_6_ biosynthesis [39], respectively; and Fur (for ferric uptake regulator), a transcription factor mainly involved in the control of iron metabolism. In the *frpB1* point mutant, the genes all displayed a similar expression pattern as well, which indicates that the downregulation of *frpB1*, resulting from the lack of ATTAAT methylation in its promoter region, is leading to the downregulation of other genes interconnected to iron metabolism (Fig. 6).

**Figure 6.**
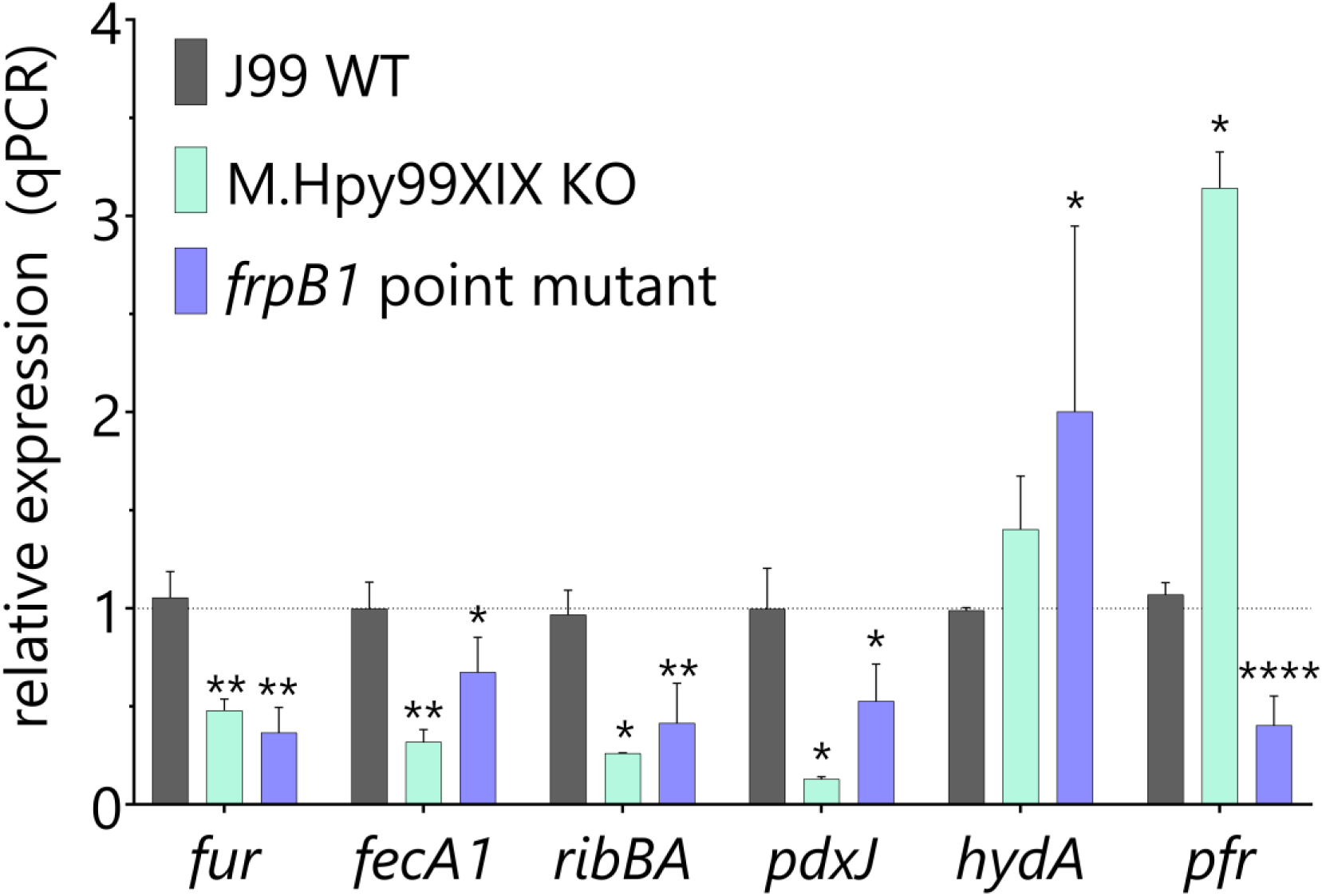
Indirect effects of ATTAAT methylation on the expression of genes related to iron metabolism. Six genes were selected based on their expression patterns in the RNA-Seq dataset and their functional role in iron homeostasis. Gene expression was quantitated by qPCR in the J99 wild-type strain, the M.Hpy99XIX knockout and the *frpB1* point mutant strain. Student’s t-test: * p<0.05, ** p<0.01, *** p<0.001, **** p<0.0001. Error bars indicate the standard deviation of 2-6 replicates.

In *H. pylori*, ferritin (Pfr) is also responsive to nickel, zinc, manganese and copper levels [36]. The discrepancy between the M.Hpy99XIX KO and the *frpB1* point mutant strains might thus be explained by general metabolic differences between strains, resulting themselves from transcriptional effects associated with different methylation profiles at other ATTAAT sites. Altogether, these results suggest that the group of differentially expressed genes observed in the M.Hpy99XIX KO strain is associated with 1) direct regulation of gene expression via promoter-bound ATTAAT methylation for a minority of genes, 2) indirect transcriptional effects mediated by core regulators and alterations of the iron metabolism resulting from the function of directly regulated genes.

### 8. M.Hpy99XIX is restricted to the *Ubiquitous H. pylori* ecospecies

We showed that the M.Hpy99XIX MTase is highly conserved in *H. pylori* and is interconnected with the regulatory pathways governing iron homeostasis, suggesting it could be important for adaptation of *H. pylori* to the stomach environment, in particular for survival under iron starvation and protection against iron toxicity. The strains in which M.Hpy99XIX is completely absent all belong to a recently described ancient group of *H. pylori*, the ‘*Hardy’* ecospecies, distinct from all other *H. pylori* strains (also termed ‘*Ubiquitous’* ecospecies) [40]. An additional analysis of the 48 *H. pylori* genomes attributed to the *Hardy* ecospecies (and isolated from humans) confirmed the complete absence of M.Hpy99XIX from this group. Interestingly, the emergence of this ecospecies is thought to be related to differences in human diets (i.e. carnivorous for *Hardy* ecospecies vs. omnivorous for *Ubiquitous*). Genetically, the *Hardy* ecospecies is characterized by 100 consistently differentiated genes from *Ubiquitous H. pylori*. Notably, differentiated genes include genes related to iron and metal homeostasis. In particular, an iron-dependent additional urease is found exclusively in *Hardy H. pylori* and several unique mutations are observed in iron uptake genes and the Fur transcriptional regulator [40].

Among both gastric and non-gastric *Helicobacter* species, orthologs of the M.Hpy99XIX MTase are only found in *H. cetorum* and *H. macacae*. In bacterial (non-*Helicobacter*) species, orthologs of M.Hpy99XIX are relatively rare, and currently found in 15 Gram-negative and 6 Gram-positive species (according to the RM systems database *REBASE*, version 410) [20]. Phylogenetically, the M.Hpy99XIX orthologs fall in two distinct clusters (Fig. 7A). The first cluster consists of genes from Gram-negative bacteria, including *H. pylori* and *H. cetorum* whereas the second cluster contains both genes from Gram-positive and Gram-negative bacteria, including *H. macacae*. Interestingly, a cognate endonuclease is observed in all organisms from the first cluster with the exception of the two *Helicobacter* species (Fig. 7B). Consequently, the loss of the cognate endonuclease in the Hpy99XIX RM system likely happened preceding its acquisition by the common ancestor of *H. pylori* and *H. cetorum*. In the second cluster, a cognate endonuclease is only observed in two species, including *H. macacae*. Altogether, the presence of an unrelated M.Hpy99XIX ortholog and a cognate endonuclease in *H. macacae* indicates that this RM system was acquired by *H. macacae* in a separate evolutionary event, unrelated to the acquisition of M.Hpy99XIX by *H. pylori*/*H. cetorum*. Furthermore, the complete absence of remnants of M.Hpy99XIX in the *Hardy* isolates suggests that this acquisition event likely took place after the divergence of the *Hardy* ecospecies from the *Ubiquitous H. pylori* isolates.

**Figure 7.**
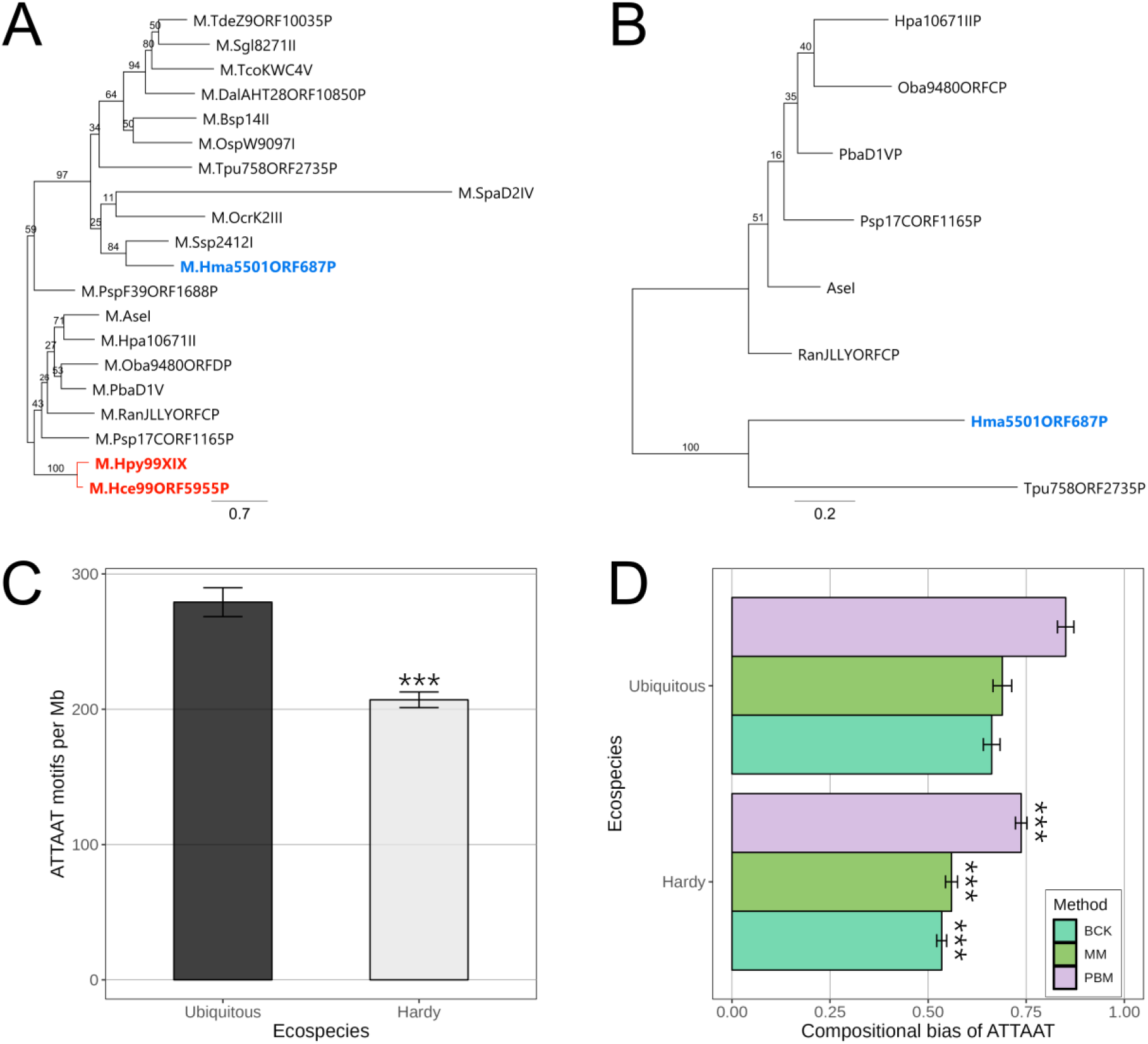
M.Hpy99XIX orthologs in bacteria and distribution of the ATTAAT motifs in *Ubiquitous* versus *Hardy* ecospecies. **A.** Phylogenetic tree of M.Hpy99XIX orthologs. **B.** Phylogenetic tree of endonucleases associated with M.Hpy99XIX orthologs. Genes from *H. pylori/H. cetorum* and *H. macacae* are highlighted in red and blue, respectively. The scale bar indicates the number of amino acid substitutions per site. **C.** Frequency of the ATTAAT motif in the *Hardy* and *Ubiquitous* ecospecies. The number of ATTAAT motifs was counted in each strain and normalized by genome size to obtain the number of motifs per million base pairs. Student’s t-test: *** p<0.001, **** p<0.00001. **D.** Compositional bias of the ATTAAT motif in the *Hardy* and *Ubiquitous* ecospecies. The compositional bias represents the ratio of observed to expected number of motifs in the genome. The expected number was calculated using three methods: a maximum order Markov chain model–based method (MM), the Pevzner and co-authors method (PBM), and the Burge and co-authors method (BCK).

We have previously identified lineage-specific variations of target motif frequency suggesting the influence of selective pressure. In this case, we asked whether the acquisition of M.Hpy99XIX in the *Ubiquitous* branch of *H. pylori* could have influenced the gain and loss of ATTAAT motifs in the genomes, possibly due to their influence on transcription. Genomes of *Hardy* strains contained on average 25% (∼ 70 motifs/Mb) fewer ATTAAT motifs than genomes of *Ubiquitous* strains (Fig. 7C). Based on an analysis of compositional bias, the ATTAAT motif is significantly more under-represented in *Hardy* than in *Ubiquitous* genomes, suggesting that the presence of the M.Hpy99XIX MTase is associated with an increased density of its target motif (Fig. 7D).

Next, we examined genes related to iron homeostasis whose expression was affected, directly or indirectly, by the M.Hpy99XIX MTase. While the *frpB1* gene was found in all *Hardy* genomes, the ATTAAT motif located within its promoter was only present in 8% of the strains (4/48). Instead, the sequences ATCAGC and ATTAGC were observed in 90% (43/48) and 2% (1/48) of the genomes. Strikingly, the *fecA1* gene was missing in 85% of the *Hardy* genomes (41/48) without any ORF remnants or traces of mobile genetic elements. In the remaining 15% of the *Hardy* genomes (7/48), an intact *fecA1* could be observed, including the promoter regions containing a single or both ATTAAT motifs in 5/7 and 2/7 cases, respectively. Other iron homeostasis genes considered in this study (*fur*, *ribBA*, *pdxJ*, *hydA* and *pfr*) were present in all *Hardy* genomes. These results indicate that the absence of the ATTAAT MTase is a conserved feature of *Hardy* strains, consistent with its role in iron homeostasis.

## Discussion

The role of bacterial orphan MTases has long remained enigmatic since, due to the lack of a cognate restriction endonuclease, they cannot serve in the defense against invading DNA. Until now, they have been typically associated with transcriptional regulation of genes involved in important cellular processes [12, 13, 41]. Deletion of specific type II MTases in *H. pylori* can lead to transcriptional changes affecting large groups of genes [12, 13]. DNA methylation has also been linked to a multitude of phenotypes, although the functional connections between (loss of) methylation and specific phenotypes have not been elucidated in most cases. For the GCGC-specific MTase M.Hpy99III, effects on specific genes have been linked to methylation of GCGC motifs within their promoter regions. However, GCGC-motifs in promoter regions were found only in a small subset of genes differentially regulated by inactivation of GCGC methylation, suggesting an important role of indirect regulation to the overall transcriptional response following disruption of GCGC methylation [12, 16].

In this study, we characterized the transcriptional response and phenotypes associated with the M.Hpy99XIX MTase and demonstrated how it is intertwined with the iron homeostasis regulatory pathway. We determined that M.Hpy99XIX is the only *H. pylori* MTase that is never associated with a cognate endonuclease and therefore the only strictly orphan MTase in the species. Other MTases have also been shown to be variably orphan in *H. pylori*. In particular, in the RM.Hpy99III RM system, targeting GCGC, the endonuclease is often truncated in *H. pylori* geographical population other than hpAfrica2 [12].

In *H. pylori*, the M.Hpy99XIX gene is present and predicted to be active in 99% of the strains. The rarity of predicted loss-of-function mutations suggests that this MTase likely serves an important function for *H. pylori*. Furthermore, strains with an inactivated M.Hpy99XIX gene might have either evolved compensatory mutations or could represent evolutionary dead-ends unable to compete with strains carrying an active M.Hpy99XIX. Strains in which the M.Hpy99XIX gene is completely missing are restricted to the recently defined *Hardy* ecospecies of *H. pylori* [40], which is distinct from the *Ubiquitous* ecospecies, the latter representing the diverse *H. pylori* strains that have been studied up to now. The absence of related M.Hpy99XIX orthologs in other *Helicobacter* species, other than *H. cetorum*, and the lack of any ORF remnants or insertion sites in *Hardy* genomes suggest that M.Hpy99XIX was never present in this group. The common ancestor of the *Hardy* and *Ubiquitous* ecospecies is thought to be extremely ancient and to have diverged before the split between the two ancestral super-lineages (the first leading to hpAfrica2 and the second to hpAfrica1 and all other populations) of *H.* pylori that happened at least 100 kya ago [42]. Therefore, it seems most likely that M.Hpy99XIX was acquired by *H. pylori Ubiquitous* after the sub-division between *Hardy* and *Ubiquitous* ecospecies, but prior to the split into the two *H. pylori* super-lineages. Interestingly, the *Hardy* ecospecies is characterized by an array of fixed mutations [40] in the Fur and ArsRS transcription factors and iron acquisition genes (*frpB4*, *tonB1*, *exbB*, *exbD*) and a specific iron-dependent urease enzyme in addition to the nickel-dependent urease already present in all *H. pylori* strains and strictly required for pathogenesis. In particular, these genetic traits are thought to represent an evolutionary strategy adapted to carnivorous diet. On the opposite, we showed that i) the promoter-bound ATTAAT motif modulating the expression of *frpB1* and ii) the entire *fecA1* gene (including its promoter with two ATTAAT motifs) are highly conserved and specific to *Ubiquitous* strains, compared to *Hardy* strains. Collectively, the acquisition of M.Hpy99XIX and the evolution of ATTA^m6^AT methylation-dependent iron-uptake genes in *Ubiquitous* strains are likely the result of synergistic evolution in response to the iron status encountered by *H. pylori* and contributed to the delineation of distinct ecospecies.

Inactivation of the M.Hpy99XIX gene led to the differential regulation of many genes. More genes were downregulated than upregulated in the absence of ATTAAT methylation, suggesting that methylation has the tendency to increase gene expression which is in line with previous observations with the M.Hpy99III MTase for instance. In a different strain, *H. pylori* P12, deletion of the M.Hpy99XIX gene also led to the differential regulation of 102 genes [13]. Around 16% of the differentially expressed genes matched between *H. pylori* J99 and P12, which is similar to the degree of strain specific variation observed with the GCGC-specific MTase M.Hpy99III in *H. pylori* strains J99 and BCM300 [12]. Furthermore, RNA-Seq experiments were performed at different optical densities (OD_600_ = 0.8 for J99 and 0.4 for P12) and analyzed using different methods (DEseq2 for J99 and TCC for P12), likely introducing additional bias into the comparison of both strains.

To understand the basis of the transcriptional effect observed in the M.Hpy99XIX KO strain, we compared all the significantly regulated genes to known *H. pylori* regulons [12, 25–30]. Interestingly, there was a substantial overlap between ATTAAT methylation-regulated genes and iron regulated genes. In *H. pylori* and other bacteria, iron homeostasis is maintained by Fur (ferric uptake regulator) and a network of differentially regulated genes according to extracellular iron availability [27, 36, 39]. In particular, the allosteric behavior of Fur produces a conformational switch when bound to a Fe^2+^ co-factor which modulates its binding to regulatory elements [24, 43, 44]. Within our dataset, we identified two genes, *fecA1* and *frpB1*, that are both involved in iron acquisition [32, 45, 46] and both carry an ATTAAT motif as well as a FUR-box in their promoter regions [35]. By specifically mutating these ATTAAT motifs, we could demonstrate that ATTAAT methylation exerts a direct transcriptional effect on these two genes. The precise mechanism behind – direct – methylation-based transcriptional regulation remains to be characterized. In *H. pylori*, a similar phenomenon was also demonstrated by targeted mutagenesis with a GCGC motif located in the promoter region of the toxin-antitoxin *jhp0832* gene, and was later validated using a reporter assay for three additional promoters [12, 16]. However, GCGC motifs within promoters tend to be located at the -13 position (relative to the transcription start site) [16] whereas ATTAAT did not appear to have any bias toward a specific position, suggesting that there might not be a common mechanism underlying methylation-based transcriptional regulation between the two MTases and that only a subset of ATTAAT motifs within promoters may have a role in gene expression depending on uncharacterized factors. Interestingly, bacterial promoters are typically AT-rich [47] and the ATTAAT motif is the only known 100% AT type II motif in *H. pylori* as well as the most over-represented in promoter regions, suggesting that the AT content of motifs likely influence their frequencies in this context.

The promoter region of the *fur* gene itself contains a FUR-box [35] but not an ATTAAT motif and thus is unlikely to be directly regulated by M.Hpy99XIX. Nevertheless, we could show that the tolerance of the M.Hpy99XIX KO strain to both high and low extracellular iron conditions was significantly decreased compared to the wild-type, indicating that M.Hpy99XIX can indirectly contribute to the maintenance of iron homeostasis in *H. pylori*. The disruption of the Fur regulatory network was further confirmed by the decreased tolerance of the M.Hpy99XIX mutants to other metal ions such as nickel or copper and to oxidative stress. Although *fur* cannot be directly regulated by M.Hpy99XIX, it was nonetheless downregulated in the absence of ATTA^m6^AT methylation, again indicating an indirect connection between both systems. Strikingly, targeted mutagenesis of the ATTAAT motif (thus preventing methylation at this site) in the promoter region of the *frpB1* gene yielded very similar phenotypes to the M.Hpy99XIX KO strain in all tested conditions. Furthermore, we could also demonstrate similar expression patterns between the *frpB1* point mutant and the M.Hpy99XIX KO strains for six genes associated with iron homeostasis, indicating that the Fur regulatory network is disrupted in both mutant strains.

Altogether, our data suggest that profound changes of the transcriptome in response to loss of a MTase can be mediated by very focused direct transcriptional regulation of a small group of genes. The effects of the M.Hpy99XIX MTase in *H. pylori* observed in this study can be summarized as a two-step model (Fig. 8). The first step consists in direct gene regulation via methylation of ATTAAT motifs located within the promoter region of two genes involved in iron acquisition, *frpB1* and *fecA1*. Changes in iron transport and thus intracellular iron content modulate the allosteric regulation of Fur [27, 36, 39], the ferric uptake regulator, preceding the second step. Switches between the apo- and holo-Fur conformations lead to either induction or repression of genes within the Fur regulatory network, including *frpB1* and *fecA1*, as well genes related to iron biosynthesis and storage [27, 36, 39, 48]. These indirect downstream effects can then generate a feedback loop and cause downstream metabolic effects such as a change in susceptibility to metal ions and tolerance to oxidative stress. In conclusion, the role of M.Hpy99XIX in gene expression and in iron homeostasis represent compelling evidence of the importance of this orphan MTase outside the confines of a typical RM system in *H. pylori* and suggest that the methylome could represent a universal and dynamic interface linking genome diversity and transcriptional regulation. Its general absence from the Hardy ecospecies is striking and will likely provide novel clues to inform future studies into the evolution and functional differentiation of the two ecospecies.

**Figure 8.**
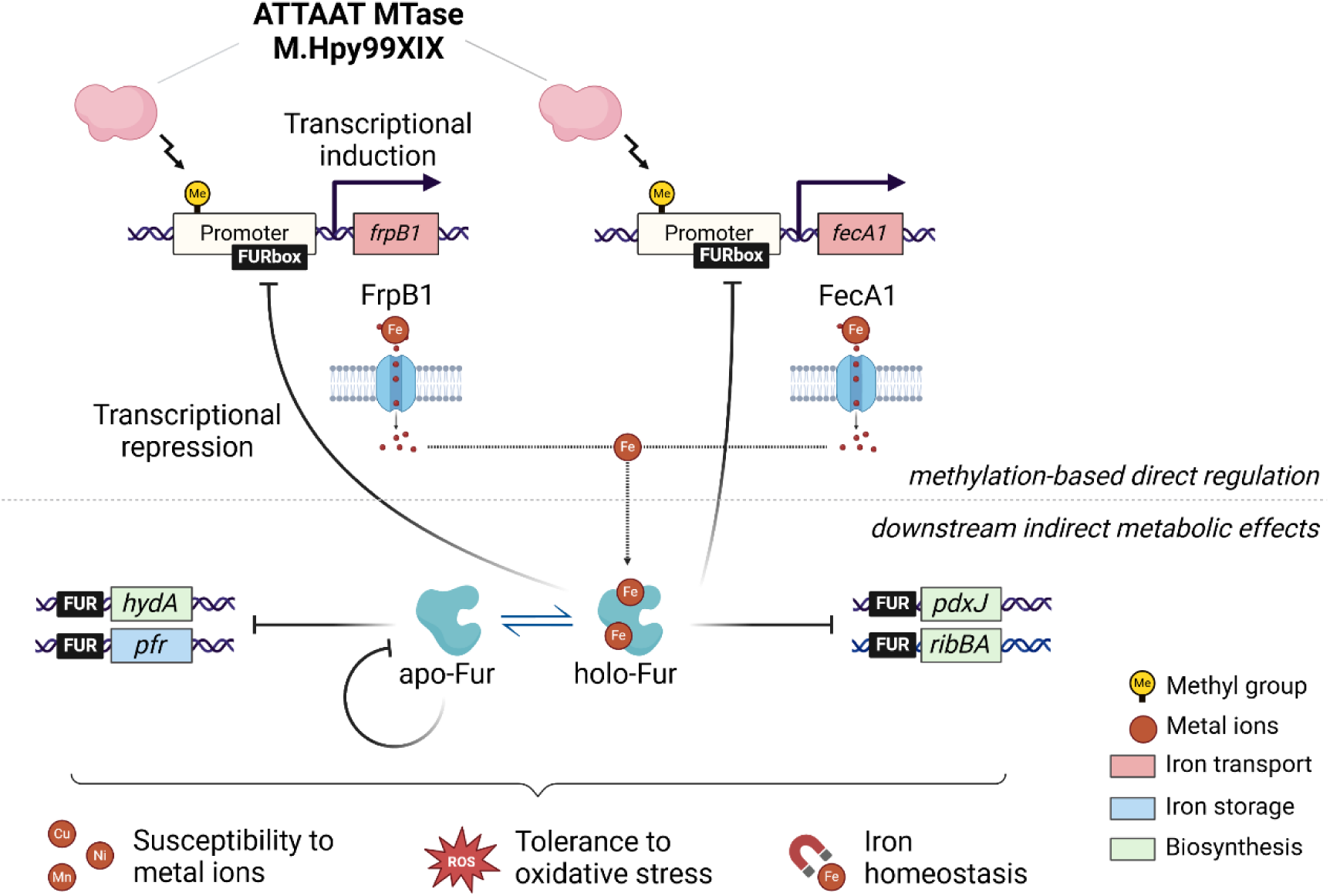
Model of the direct and indirect transcriptional and phenotypic effects of the M.Hpy99XIX methyltransferase. The effects of ATTAAT methylation by M.Hpy99XIX can be described in two steps. The first step consists in direct gene regulation via methylation of an ATTAAT motif within the promoter region, demonstrated by qPCR and RNA-Seq for two genes involved in iron acquisition, *frpB1* and *fecA1*. Changes in iron acquisition and intracellular iron content influence the allosteric regulation of Fur, the ferric uptake regulator, preceding the second step. Switches between the apo- and holo-Fur conformations lead to both induction or repression of genes within the Fur regulatory network, including *frpB1* and *fecA1* as well genes related to iron biosynthesis and storage, as shown by qPCR. These indirect downstream effects can then generate a feedback loop and cause downstream metabolic effects such a changes in susceptibility to metal ions and tolerance to oxidative stress, as demonstrated by phenotype growth assays in multiple environmental conditions.

## Methods

### Analyses of type II RM systems frequencies in *H. pylori* and other bacterial species

The genome collection used to determine the frequency of methyltransferases and endonucleases, normalized across four geographical populations of *H. pylori* (hpAfrica1, hpAfrica2, hpEurope and hpAsia2) is based on our previous analysis of the methylome diversity in *H. pylori* [8]. The genome collection used to characterize the absence and mechanisms of inactivation of the M.Hpy99XIX MTase was built out of 1005 complete genomes from the HpGP project [21] as well as 24 additional complete genomes from several *H. pylori* evolutionary studies [49–51] and reference strains [5, 52–55]. MTase and endonuclease sequences from *H. pylori* were retrieved from the Rebase database [20] and their presence in the collection was assessed using BLASTn and BLASTp [56]. To infer the location of transcription start sites and the sequence of promoter regions, previously published promoter regions of 50 bp upstream sequence of the TSS obtained using the dRNA-seq method [57] were mapped to the genome collection using bbmap [58] with the following settings: vslow, k=8, maxindel=200, minid=0.5, secondary=f and ordered=t.

Sequences of orthologs of the M.Hpy99XIX MTase and endonucleases were obtained from the RM system database *REbase* [20]. Protein sequences were aligned using MAFFT [59] and phylogenetic trees were built using RAxML with the rapid hill-climbing mode, the GAMMA LG model and 100 bootstrap replicates [60].

### Analyses of target motifs frequencies and distribution across the *H. pylori* genome

Frequency and position of MTase target motifs were analyzed using the Biostrings 2.58.0 Bioconductor R package [61]. Fold-change of the target motifs in promoter regions was calculated as the log2 of the ratio of motif frequency within promoter over motif frequency in the genome. A positive fold-change indicates over-representation of the motif in promoters whereas a negative fold-changed indicates under-representation. The position of the ATTAAT and TATAAT motif across all promoter regions was summarized by computing a probability density function using the R package mclust [62].

Enrichment of the ATTAAT motif in coding sequences was calculated using the distAMo software [63]. A z-score of 2 and -2 was considered as significant enrichment and depletion, respectively.

Compositional bias of the ATTAAT motif was calculated using sing the methods of Burge and co-authors [64], Pevzner and co-authors [65], and a maximum-order Markov chain model implemented in the CBcalc software [66].

### Bacterial culture and growth curves

*H. pylori* strains were cultured from frozen stocks on blood agar plates or in liquid cultures as described previously [12]. Liquid cultures of bacteria were grown in brain heart infusion broth (BHI, BD Difco, Heidelberg, Germany) with yeast extract (2.5 g l^-1^), 10% heat-inactivated horse serum and a mix of antibiotics (vancomycin (10 mg l^-1^), polymyxin B (3.2 mg l^-1^), amphotericin B (4 mg l^-1^) and trimethoprim (5 mg l^-1^)). Anaerocult C gas producing bags (Merck, Darmstadt, Germany) were used to generate microaerobic conditions in airtight jars (Oxoid, Wesel, Germany). Growth curves were generated for the different strains of 10 mL liquid culture starting at an OD_600_ of 0.06 in 50 mL falcons, incubated with shaking (140 rpm, 37 °C, microaerobic conditions) and measured after 8, 24, 32, 48, 56, 72 and 80 hours of growth.

### RNA extraction

For RNA extraction, 5 mL of bacterial culture grown in liquid media to OD_600_ of 0.8 was pelleted (4°C, 6000 x G, 5 min), snap-frozen in liquid nitrogen and stored at -80 °C. Afterwards, bacterial pellets were disrupted with a Precellys 24 homogenisator (Bertin Technologies, Montigny-le-Bretonneux, France) using Lysing Matrix B 2 mL tubes containing 0.1 mm silica beads (MP Biomedicals, Eschwege, Germany). Isolation of RNA was performed with the RNeasy Mini Kit (Qiagen, Hilden, Germany) with an on-column digestion DNase digestion with DNase I and a second DNase treatment using the TURBO DNA-free Kit (Invitrogen, Carlsbad, United States). Isolated RNA was checked for absence of DNA contamination by PCR of the 16S rRNA gene [12]. The RNA concentration was measured using the NanoDrop 2000 spectrophotometer (Peqlab Biotechnologies, Erlangen, Germany). The quality of the RNA was tested with the 4200 TapeStation System using RNA Screen Tapes (Agilent, Waldbronn, Germany). All RINe numbers of prepared RNA were higher than 8.8, confirming high quality and little RNA degradation.

### Transcriptomic analysis: RNA sequencing

RNAseq was performed on an Illumina sequencing platform (2×150 bp, >10M read pairs) by Eurofins (Ebersberg, Germany). Probe-based ribosomal RNA depletion was performed for each sample prior to mRNA fragmentation, strand-specific cDNA synthesis and library preparation. Three biological replicates were used for each condition. Data was analyzed with R package DESeq2 [67]. An absolute fold change larger 1 and an FDR adjusted p-value below 0.01 were used as a cutoff.

### Construction of *H. pylori* mutant strains

Inactivation of the MTase was carried out by the Multiplex Genome editing (MuGent) technique as previously described [68, 69]. Briefly, we use natural co-transformation with a chloramphenicol resistance cassette within the non-essential *rdxA* locus as a selective marker in addition to a non-selective PCR product carrying a construct of interest. To generate gene knockouts, 1 kb regions up- and downstream of the MTase were amplified by PCR, digested and ligated into a digested pUC19 vector. A PCR product of the deletion construct was then used as a non-selective product in the co-transformation reaction. To perform targeted mutagenesis and generate point mutants, a 1 kb fragment of the target sequence was cloned into the pUC19 vector and an inverse PCR was performed using primers carrying the desired mutation. The resulting PCR product was then digested with DpnI (NEB, Frankfurt am Main, Germany), PCR purified with the QIAquick PCR Purification Kit (Qiagen, Hilden, Germany) and re-ligated using the Quick Ligation Kit (NEB, Frankfurt am Main, Germany) before transformation into *E. coli* MC1061 [12, 70]. PCR products of target sequence were then used as a non-selective product in the co-transformation reaction [68, 69]. Sanger sequencing was used to verify the acquisition of the desired mutations.

Lastly, we generated a functional complementation of the MTase gene in the J99 MTase KO strain using the pADC/CAT suicide plasmid approach, as described previously [12, 71]. The MTase gene was inserted into the urease locus under control of the urease operon promoter.

The *H. pylori* mutants were checked via PCR and selected on antibiotic-containing plates. The absence or recovery of ATTAAT methylation was checked by digestion of gDNA with AseI (NEB, Frankfurt am Main, Germany). Primers and plasmids used in this study are described in Table S1.

### Transcriptomic analysis: qPCR

One µg of RNA was used for cDNA synthesis using the SuperScript^TM^ III Reverse Transcriptase (Thermo Fischer Scientific, Darmstadt, Germany) as described previously [72]. qPCR was performed with gene specific primers (Table S1) and SYBR Green Master Mix (Qiagen, Hilden, Germany). Reactions were run in a BioRad CFX96 system. Standard curves and samples were run as biological quadruplicates and technical duplicates. Samples were normalized to an internal 16S rRNA control for quantitative comparison. The quality standards used correspond to those in our previously published MIQE [12].

### Growth assays

Susceptibility to iron (1 mM (NH4)_2_Fe(SO_4_)_2_), copper (0.35 mM CuSO_4_), zinc (0.5 mM ZnCl_2_), manganese (0.5 mM MnCl_2)_, nickel (20 µM NiSO₄), oxidative stress (1 µM paraquat) and iron chelation (75 µM dpp (2,2′-dipyridyl)) was tested by adding the respective compounds to 100 µL liquid cultures grown in 96-well plates with a starting OD_600_ of 0.1. The OD_600_ was measured for the following 18 h with a CLARIOstar microplate reader (BMG Labtech, Ortenberg, Germany) under constant 10 % CO_2_. The average of the last 10 measurements obtained over a 20-minute period were used as the end-point OD_600_ for each measurement. The end-point values obtained for each compound were normalized to the final OD_600_ of an untreated control.

## Reproducibility

All data was analyzed using R version 4.2.3, Geneious 11.0.4 or GraphPad Prism version 9.4.1.

## Declaration of competing interest

The authors declare they have no competing interests.

## Acknowledgements

We thank Gudrun Pfaffinger and Evelyn Weiss for their excellent technical assistance.

## Author contributions

S.S. and F. A. designed the study. W.G. and F. A. performed the experiments and bioinformatic analysis. All authors interpreted the results, wrote and revised the manuscript, and approved the final version.

## Notes

### Competing Interest Statement

The authors have declared no competing interest.

