## Supplemental Figures S1-S8 for "The *Helicobacter pylori* orphan ATTAAT-specific methyltransferase M.Hpy99XIX plays a central role in the coordinated regulation of genes involved in iron metabolism"

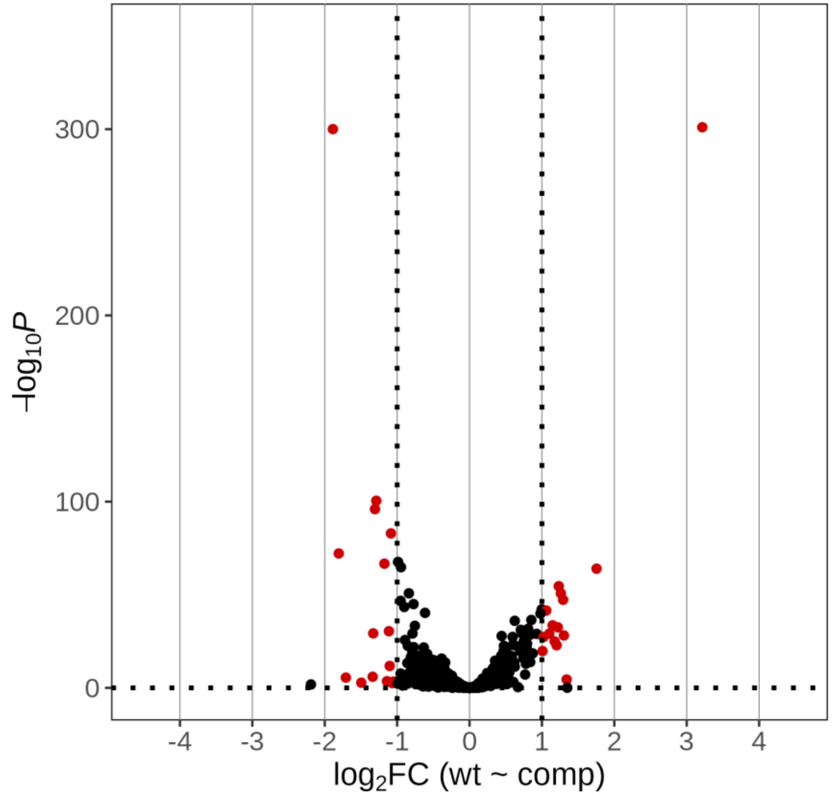


**Figure S1. Transcriptomic effect of M.Hpy99XIX complementation in *H. pylori* strain J99.** Volcano plot of *H. pylori* J99 wild-type compared to *H. pylori* J99 M.Hpy99XIX complementation strain. A negative and positive $Log_2$ fold change indicate a gene downregulated or upregulated in the complementation strain compared to the wild-type, respectively. Genes with an adjusted p-value $< 0.01$ and an absolute $log_2$ fold change $> 1$ are shown in red.

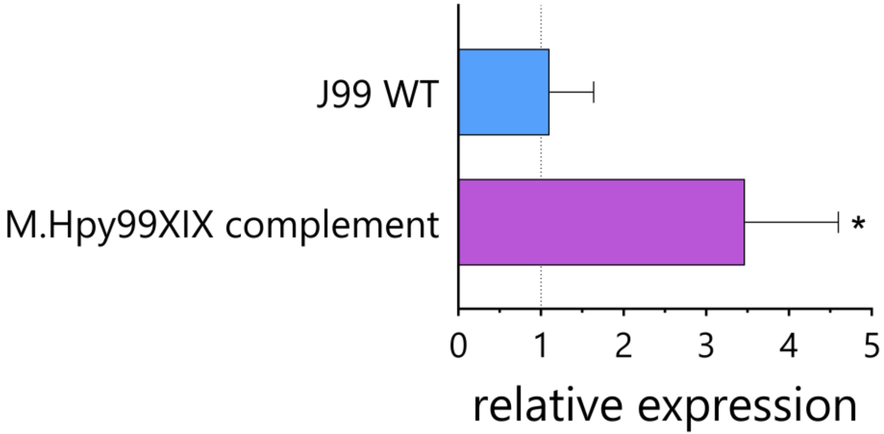


**Figure S2. Expression of the M.Hpy99XIX methyltransferase in the complementation mutant.** qPCR analysis of the expression of M.Hpy99XIX in the complementation mutant compared to the J99 WT. Student's t-test: * $p<0.05$. Error bars indicate the standard-deviation of 2-3 replicates.

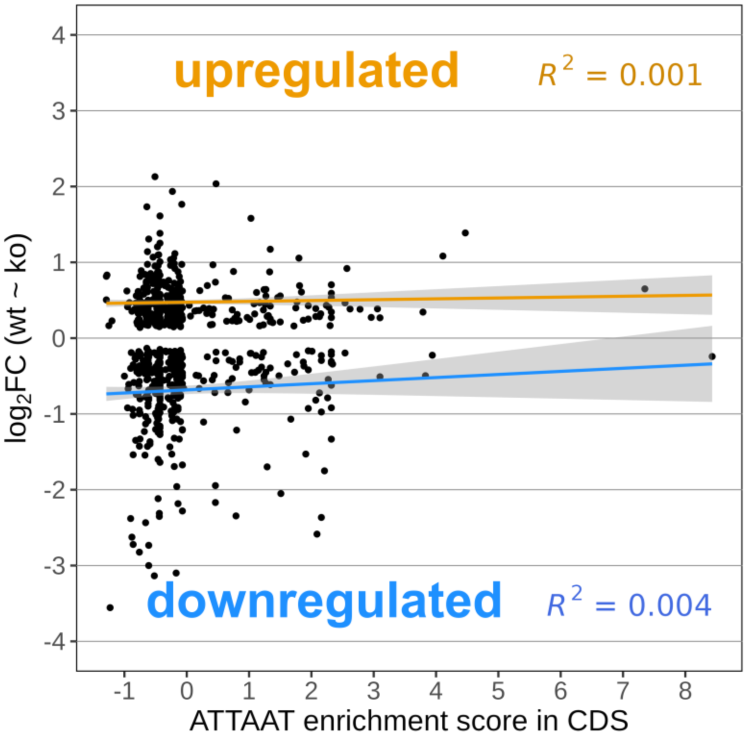


**Figure S3. Association of ATTAAT motifs located in coding sequences (CDS) with transcriptomic changes in the M.Hpy99XIX KO strain.** Enrichment score of the ATTAAT motif in CDS (calculated using the DistAMo algorithm, see Methods) is plotted on the x-axis again the $Log_2FC$ of differentially expressed genes (J99 WT versus M.Hpy99XIX KO) on the y-axis. A linear regression was performed separately for up-regulated genes ($Log_2FC > 0$, orange line) and downregulated genes ($Log_2FC < 0$, blue line). Confidence intervals are shown as a grey area for each regression. The coefficient of determinations (R-squared) are indicated for each regression and, in both cases, suggest an absence of correlation between the enrichment of the ATTAAT motifs in CDS and the transcriptomic changes observed in the M.Hpy99XIX KO strain.

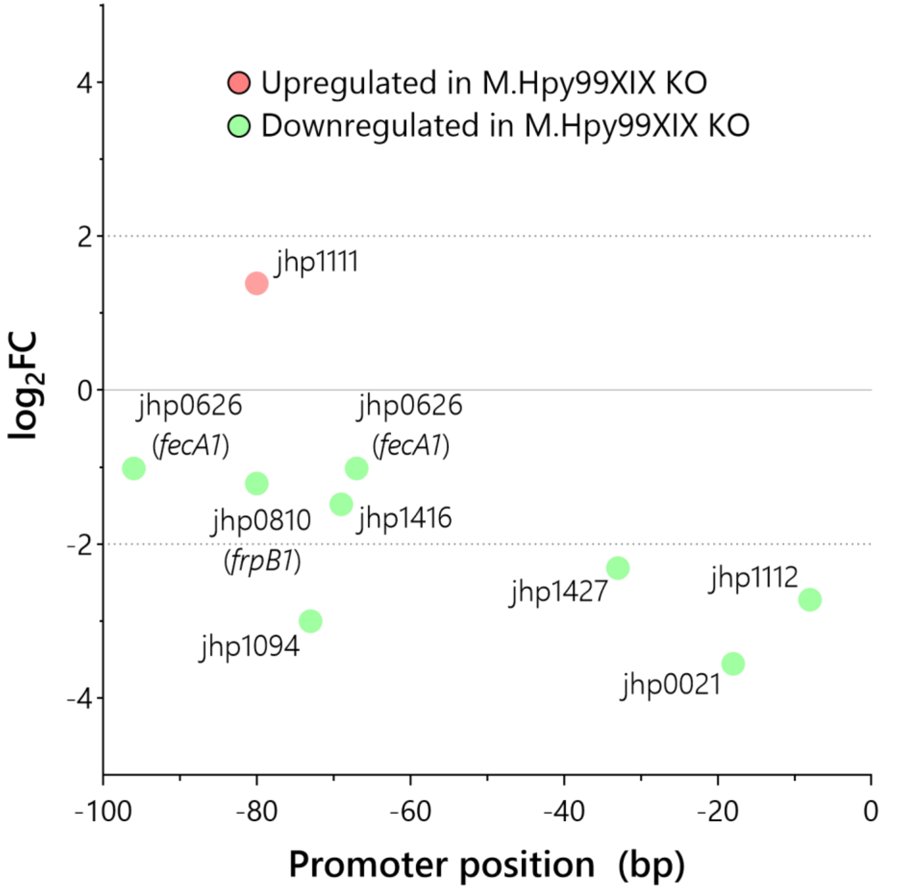


**Figure S4. Characterization of the motif position in promoter regions of differentially expressed genes.** The position of the first ATTAAT motif in the promoter region defined as 100 bp upstream of the TSS is displayed for the 8 differentially expressed genes according to their $\log_2$ fold change.

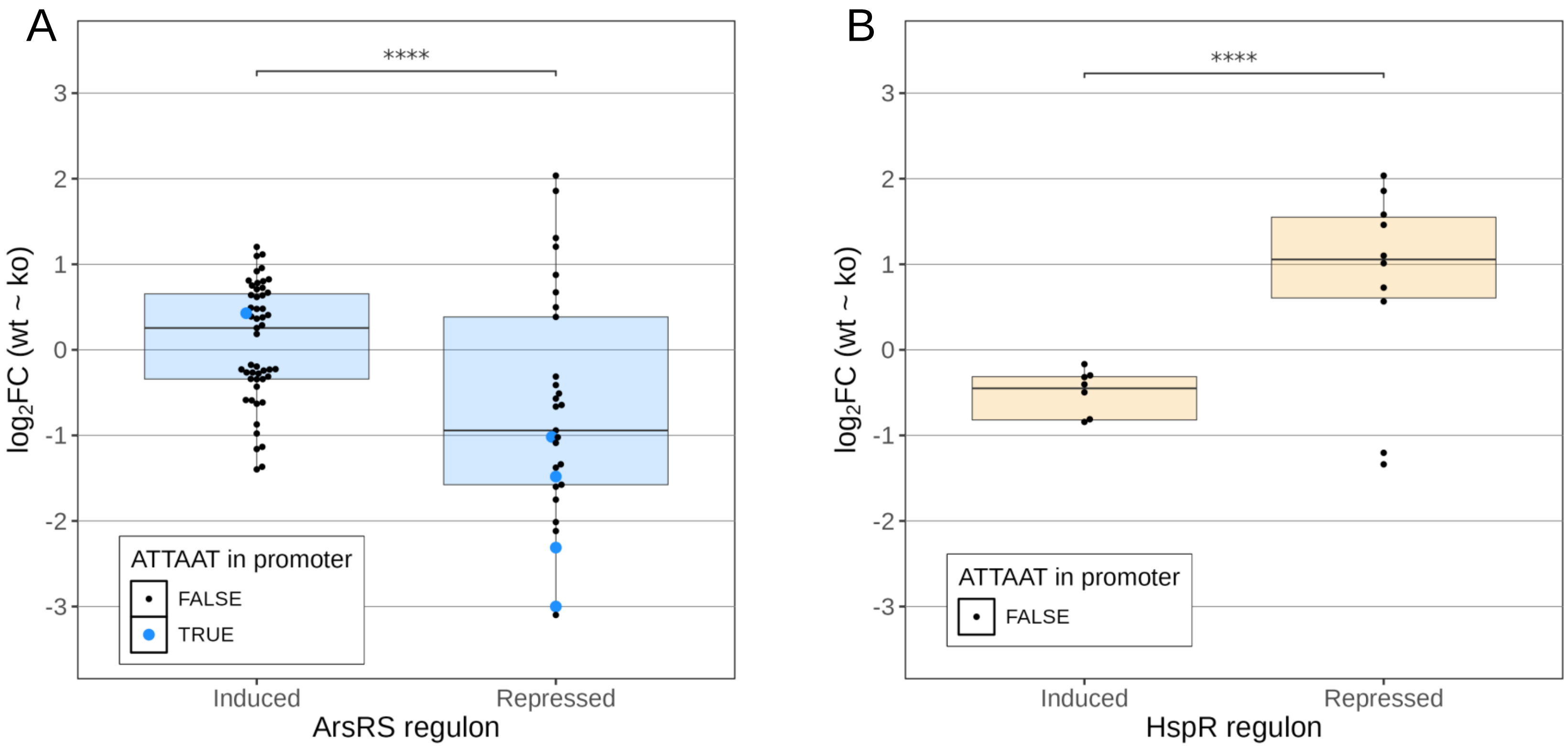


**Figure S5. Comparison between M.Hpy99XIX transcriptional effects and environment responsive regulons. A.** Comparison of RNA-Seq expression data of M.Hpy99XIX KO against WT with the ArsRS-responsive regulon (Loh et al. 2021) split into induced and repressed genes. Student's t-test: *** p<0.001, **** p<0.00001. **B.** Comparison of RNA-Seq expression data of M.Hpy99XIX KO against WT with the HspR-responsive regulon (Roncarati et al. 2007) split into induced and repressed genes. Student's t-test: *** p<0.001, **** p<0.00001. $Log_2$ fold-changes of significantly differentially expressed genes (adjusted p-value < 0.01) are represented by individual points and by a boxplot. The presence of an ATTAAT motif within -100 bp of the TSS of each gene is indicated by a larger coloured point.

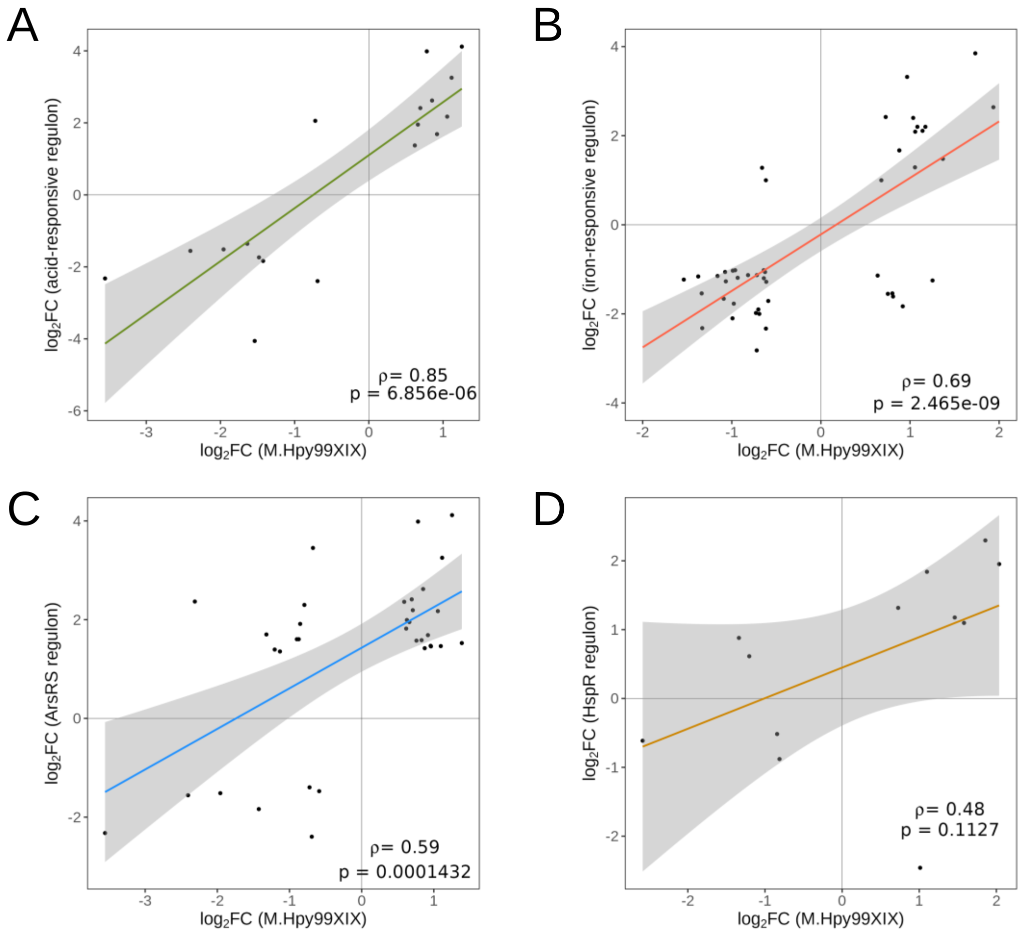


**Figure S6. Association between M.Hpy99XIX transcriptional effects and environment responsive regulons.** Associations between genes with >|50%| differential expression (|$\log_2$ fold-change| > $\log_2(1.5)$ and p.adj < 0.01) observed in the M.Hpy99XIX KO strain and in different regulons were calculated using the Pearson's correlation coefficient (indicated by $\rho$ and p-value indicated below). A linear regression with 95% confidence interval is also shown. **A.** Acid-responsive regulon. **B.** Iron-responsive regulon. **C.** ArsRS regulon. **D.** HspR regulon.

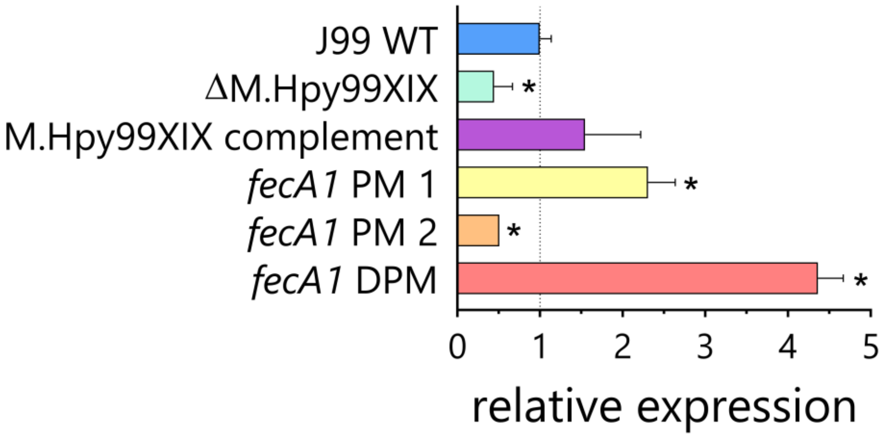


**Figure S7. Characterization of the effect of ATTAAT methylation within the promoter region of *fecA1.*** Comparison of *fecA1* expression in the M.Hpy99XIX knockout, the point-mutants (PM) and double point-mutant (DPM) to the wild-type strain by qPCR. Student's t-test: * $p<0.05$. Error bars indicate the standard-deviation of 2-3 replicates.

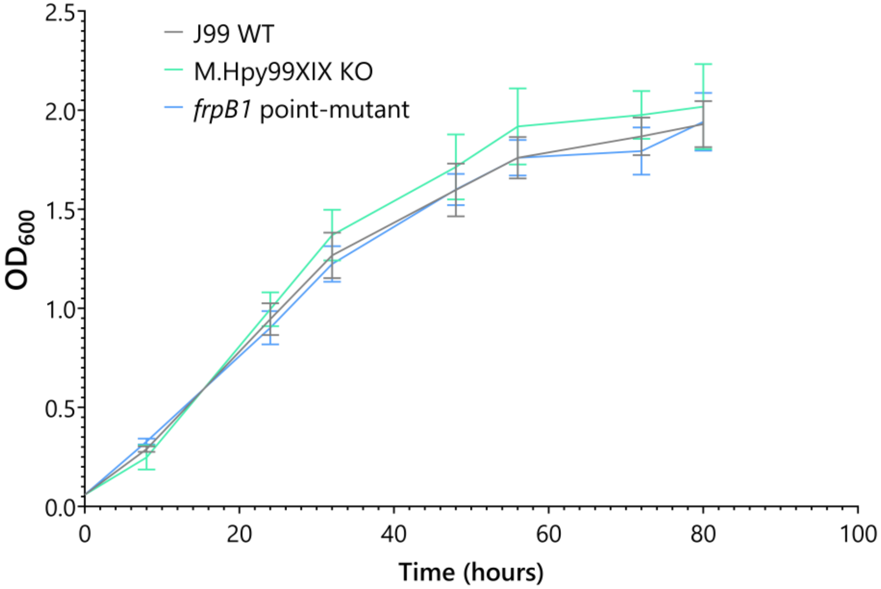


**Figure S8. Characterization of the effect of global (M.Hpy99XIX KO) and local (*frpB1* point mutant) ATTAAT methylation on growth of *H. pylori* strain J99.** Comparison of growth in liquid culture between the M.Hpy99XIX knockout and the *frpB1* point mutant over a time course of 80 hours. Error bars indicate the standard deviation of 5 replicates.
